# Identifying extrinsic versus intrinsic drivers of variation in cell behaviour in human iPS cell lines from healthy donors

**DOI:** 10.1101/285627

**Authors:** Alessandra Vigilante, Anna Laddach, Nathalie Moens, Ruta Meleckyte, Andreas Leha, Arsham Ghahramani, Oliver J. Culley, Annie Kathuria, Chloe Hurling, Alice Vickers, Erika Wiseman, Mukul Tewary, Peter Zandstra, HipSci Consortium, Richard Durbin, Franca Fraternali, Oliver Stegle, Ewan Birney, Nicholas M. Luscombe, Davide Danovi, Fiona M. Watt

**Affiliations:** Centre for Stem Cells and Regenerative Medicine, King’s College London, Floor 28, Tower Wing, Guy’s Hospital, Great Maze Pond, London, SE1 9RT, UK; European Molecular Biology Laboratory, European Bioinformatics Institute, Wellcome Trust Genome Campus, Hinxton, Cambridge, CB10 1SA, UK; The Wellcome Trust Sanger Institute, Wellcome Trust Genome Campus, Hinxton CB10 1SA, UK; The Francis Crick Institute, 1 Midland Road, London NW1 1AT; Randall Division, King’s College London, New Hunts House, Great Maze Pond, London SE1 9RT, UK; The Donnelly Centre for Cellular and Biomolecular Research, University of Toronto, 160 College Street, Room 1116, Toronto, Ontario, Canada M5S 3E1; UCL Genetics Institute, Department of Genetics, Evolution and Environment, University College London, Gower Street, London WC1E 6BT, UK

**Author notes:** Present address: GlaxoSmithKlein, Gunnels Wood Road, Stevenage, Herts, SG1 2NY, UK. Present address: Sobell Department, University College London Institute of Neurology, Queen Square House, Queen Square, London WC1N 3GB, UK. Present address: University Medical Center Göttingen, Georg-August-Universität, Department of Medical Statistics, Humboldtallee 32, 37073 Göttingen, Germany (Andreas Leha).

## Abstract

Large cohorts of human induced pluripotent stem cells (iPSCs) from healthy donors are a potentially powerful tool for investigating the relationship between genetic variants and cellular phenotypes. Here we integrate high content imaging, gene expression and DNA sequence datasets from over 100 human iPSC lines to explore the genetic basis of inter-individual variability in cell behaviour. By applying a dimensionality reduction approach, Probabilistic Estimation of Expression Residuals (PEER), we extracted factors that captured the effects of intrinsic (genetic) and extrinsic (environmental) conditions. We identified genes that correlated in expression with intrinsic and extrinsic PEER factors and mapped outlier cell behaviour to expression of genes containing rare deleterious SNVs. Our study thus establishes a strategy for determining the genetic basis of inter-individual variability in cell behaviour.

## Introduction

Now that the applications of human induced pluripotent stem cells (hiPSC) for disease modelling and drug discovery are well established, attention is turning to the creation of large cohorts of hiPSC from healthy donors. These offer a unique opportunity to examine common genetic variants and their effects on gene expression and cellular phenotypes^1–5^. Genome-wide association studies (GWAS) and quantitative trait locus (QTL) studies can be used to correlate single nucleotide polymorphisms (SNPs) and other genetic variants with quantitative phenotypes^6^. As part of this effort, we recently described the generation and characterisation of over 700 open access hiPSC lines derived from 301 healthy donors as part of the Human Induced Pluripotent Stem Cells Initiative (HipSci)^5,7^. In addition to creating a comprehensive reference map of common regulatory variants affecting the transcriptome of hiPSC, we performed quantitative assays of cell morphology and demonstrated a donor contribution in the range of 8–23% to the observed variation. In the present study we set out to identify causative genetic variants.

Previous attempts using lymphoblastoid cell lines to link genetics to in vitro phenotypes have had limited success^8,9^. In that context, confounding effects included EBV viral transformation, the small number of lines analysed, variable cell culture conditions and line-to-line variation in proliferation rate. These non-genetic factors decrease the power to detect true relationships between DNA variation and cellular traits^8^. In contrast, we have access to a large number of hiPSC lines from healthy volunteers, including multiple lines from the same donor. In addition, HipSci lines present a substantially lower number of genetic aberrations than reported for previous collections^5,10,11^. Cells are examined at low passage number, and cell properties are evaluated at single cell resolution during a short time frame, using high throughput quantitative readouts of cell behaviour.

Stem cell behaviour reflects both the intrinsic state of the cell^12,13^ and the extrinsic signals it receives from its local microenvironment, or niche^14,15^. We hypothesised that subjecting cells to different environmental stimuli increases the likelihood of uncovering links between genotype and cell behaviour. For that reason, we seeded cells on different concentrations of the extracellular matrix (ECM) protein fibronectin that support cell spreading to differing extents, and assayed the behaviour of single cells and cells in contact with their neighbours. We took a ‘cell observatory’ approach, using high-throughput, high content imaging to gather data from millions of cells 24 hours after seeding. We then used a multidimensional reduction method, Probabilistic Estimation of Expression Residuals (PEER)^16^ to reveal underlying structure in the dataset, and correlated cell behaviour with expression of a subset of genes and the presence of rare deleterious SNVs. The strategy we have developed bridges the gap between genetic and transcript variation on the one hand and cell phenotype on the other, and should be of widespread utility in exploring the genetic basis of interindividual variability in cell behaviour.

## Results

### Generation and characterisation of the lines

We analysed 110 cell lines from the HipSci resource^2^ (Supplementary Table 1). Of these, 99 lines were reprogrammed by Sendai virus and 11 using episomal vectors. 100 lines came from 65 healthy research volunteers; thus several lines were derived from different clones from the same donor. Seven lines came from 7 individuals with Bardet-Biedl Syndrome and 3 were non-HipSci control lines. Out of the total, 102 of the lines were derived from skin fibroblasts, 2 from hair follicles and 6 from peripheral blood monocytes. All lines were subjected to the quality controls specified within the HipSci production pipeline, including high PluriTest (Stem Cell Assays) scores and the ability to differentiate along the three embryonic germ layers. All the cell lines were reprogrammed on feeders and all but 6 lines were cultured on feeders prior to phenotypic analysis (Supplementary Table 1). Cells were examined between passages 15 and 45.

### Cell behaviour assays

To quantitate cell behaviour at single cell resolution we used the high-content imaging platform that we described previously^17^. Cells were disaggregated and resuspended in the presence of 10 μM Rho-associated protein kinase (ROCK) inhibitor to minimise cell clumping. In order to vary the extrinsic conditions for cell adhesion and spreading, cells were seeded on 96 well plates coated with 3 different concentrations of fibronectin – 1, 5 and 25 μg/ml (Fn1, Fn5, Fn25), with Fn1 representing a suboptimal concentration for cell attachment and spreading. After 24h of culture in the presence of ROCK inhibitor, cells were labelled with EdU for 30 min (to detect proliferative cells), fixed, and stained with DAPI (to visualise nuclei) and CellMask (to visualise cytoplasm) (Figure 1A). Under these conditions, over 95% of cells were in the pluripotent state, as evaluated by Oct4 labelling.

**Figure. 1.**
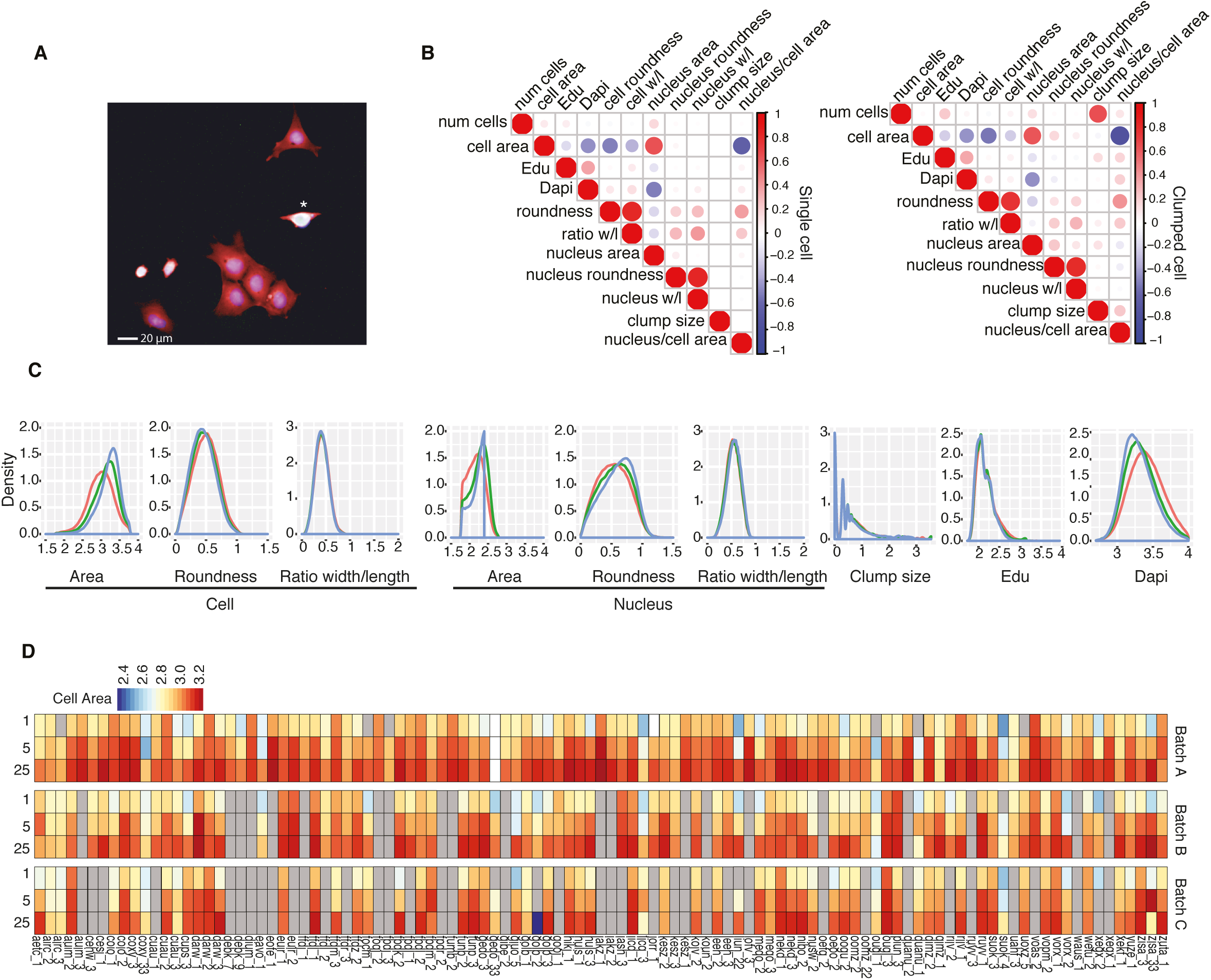
Description of phenotypic dataset. **(A)** Microscopic image showing cells 24h after plating. Red: cell mask (cytoplasm); white: EdU incorporation (DNA synthesis, one EdU+ cell marked with asterisk); blue: DAPI (nuclei). Scale bar: 20 μm**. (B)** Correlation of different phenotypic measurements in single (left) and clumped (right) cells. **(C)** Distribution of main phenotypic features of all cell lines on three fibronectin concentrations (Fn1, red; Fn5, green; Fn25, blue). Y axis: density. **(D)** Heatmap of mean cell area measurements for each cell line on three fibronectin concentrations in three independent experiments. Grey boxes correspond to data unavailable or not satisfying the minimum number of cells per well.

Three replicate wells were seeded per cell line and cell lines were analysed in up to three independent experiments. Wells containing technical triplicates of each fibronectin concentration were randomised per column *(e.g*. 1–5–25; 5–25–1) to obviate edge and position effects. Technical replicates of the same cell line were randomised in rows and one line, previously reported as A1ATD-iPSC patient 1^18^, was included to control for biological variation between experiments.

We extracted a total of 11 measurements, 10 per cell *(i.e*. object) plus the number of cells per well for approximately 2 million cells (Figure 1B-C). Cell features included the derived nucleus/cell area and clump size, a context feature allowing us to distinguish between cells that had attached as individuals and cells in groups (Figure 1B-C). Some features were positively correlated with one another, such as cell area and nuclear area, while in other cases, such as cell area and cell roundness, there was an inverse correlation (Figure 1B). The phenotypic features were processed as described previously^17^, *i.e*. well-based measurements were normalised in value (log10 or square transformation) and aggregated across the cells in each well by taking the average and standard deviation. For EdU incorporation, median pixel intensity raw values per cell were used to extract a well-based measure of the fraction of EdU positive cells^17^. This resulted in a final list of 52 features including clumps (Supplementary Table 2).

The scale and complexity of the cell phenotype dataset is illustrated in Figure 1D, in which the mean value of cell area is represented for all cell lines, for three fibronectin concentrations and three biological replicates. This highlights the variance we observed between replicate experiments. It also shows a consistent effect of fibronectin concentration, with cells exhibiting greater cell area on the highest concentration (see also Figure 1C). It reveals some variability for cell lines derived from the same donor, denoted by a common 4 letter code (Figure 1D). Similar results were obtained for other raw phenotypic features (Supplementary Figure 2).

**Figure 2.**
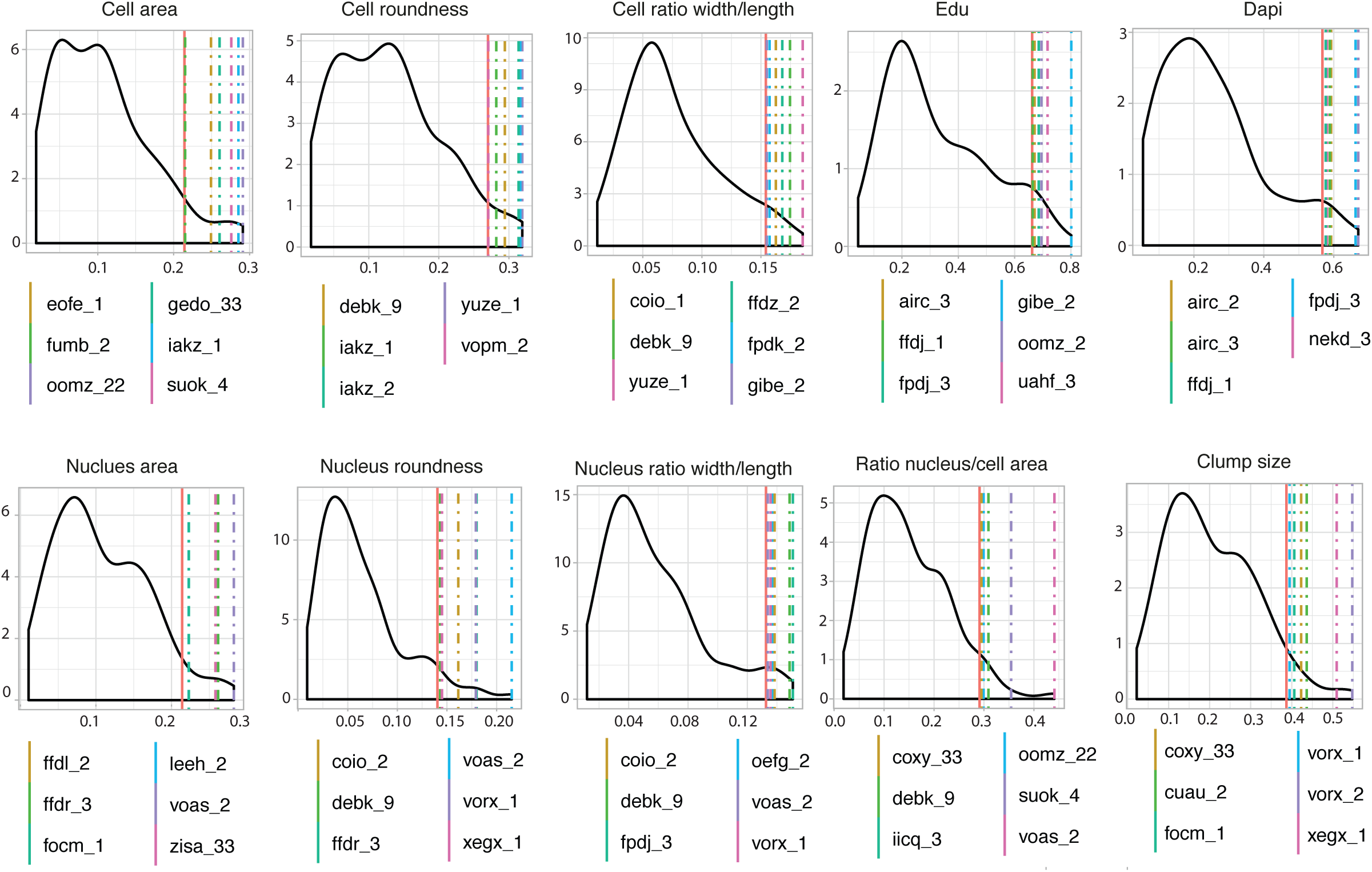
Identification of outlier cell lines for individual phenotypes. The distribution of the Kolmogorov-Smirnov statistic (D) obtained by performing the Kolmogorov-Smirnov test of the distributions of each raw phenotypic feature for each individual cell line compared to all cell lines. 95^th^ percentile threshold is shown as a red line together with values of individual outlier lines (colour coded).

### Identification of outlier cell lines

We next identified outlier cell lines, defined as lines that deviated significantly from modal phenotypic values. To do this we performed a Kolmogorov–Smirnov test of the distributions of each raw phenotypic feature for each individual cell line compared to all cell lines (Figure 2) and defined outliers as cell lines with a statistic (D) value above the 95th percentile. Out of the 110 lines analysed, 36 lines from 30 donors exhibited outlier behaviour for one or more phenotypic feature (Figure 2).

In support of a genetic contribution to outlier cell behaviour, in several cases two independent lines from the same donor exhibited the same outlier behaviour. For example iakz_1 and iakz_2 were outliers for cell roundness, while airc_2 and airc_3 were outliers for DAPI nuclear staining intensity. In addition, where two phenotypes were positively or negatively correlated *(e.g*. cell area and cell roundness) some cell lines were outliers in both categories *(e.g*. iakz_1).

### Contribution of intrinsic and extrinsic factors to variation in cell phenotypes

In order to explore how extrinsic (i.e. different fibronectin concentrations), intrinsic (i.e. cell line or donor specific) and technical or biological components (covariates) contributed to the observed variation in cell phenotypes, we applied a dimensionality reduction approach called Probabilistic Estimation of Expression Residuals (PEER)^16^. PEER was originally implemented for gene expression data and to our knowledge has not been applied previously to multidimensional reduction of phenotypic data. It is a collection of Bayesian approaches that takes as input measurements (i.e. expression value) and covariates and then extract factors that explain hidden portions of the variability. This is fundamental because unobserved, hidden factors, such as cell culture conditions can have an influence on large numbers of cells and many studies have demonstrated the importance of accounting for hidden factors to achieve a stronger statistical discrimination signal^54 55 56^.

In our analysis, PEER takes the 52 phenotypic measurements and covariates *(i.e*. fibronectin concentration, experimental replicates, individual donors) as inputs and outputs hidden factors that explain portions of the variance and can be treated as new synthetic phenotypes. Using the automatic relevance detection (ARD) parameters to understand which dimensions are needed to model the variation in the data^16^, the optimal number of PEER factors was set to 9 (Supplementary Figure 3). For a single factor (PEER factor 1) the effect of fibronectin concentration was apparent and significant between each condition (Figure 3A and Supplementary Figure 4). This factor, which we named the ‘extrinsic factor’, accounted for ~30% of the total variance (Figure 3A, C, E and Supplementary Figure 3). This contrasted with Principal Component Analysis for which the effects of Fibronectin concentrations was apparent on both PC1 (27.7%) and PC2 (15.3%). Figure 3E shows that the mean of the total number of attached cells, cell and nucleus area and DAPI staining intensity have the largest effect onto the extrinsic factor.

**Figure 3.**
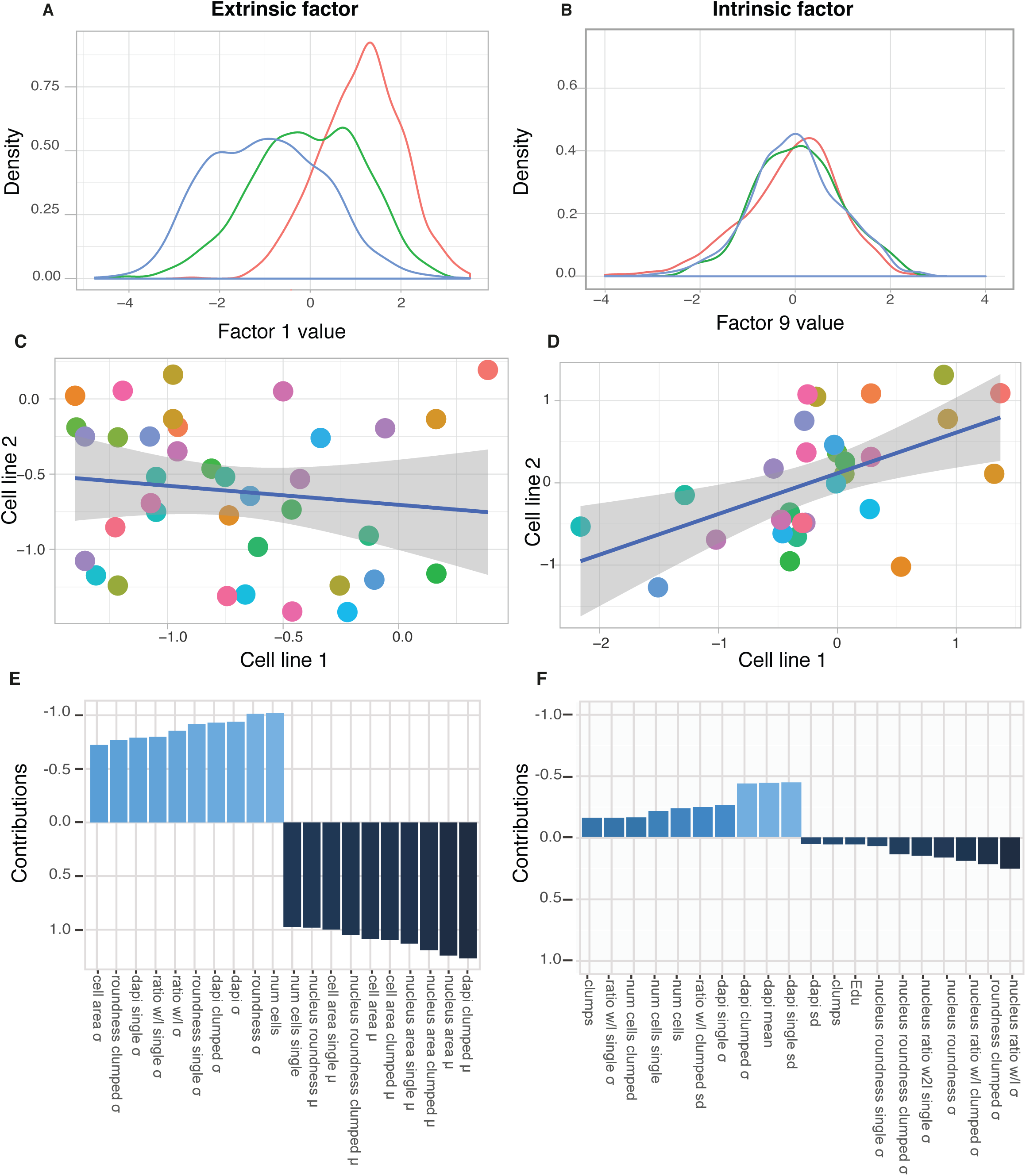
Synthetic phenotypic features capture extrinsic and intrinsic contributions to variance. **(A-B-C-D)** Plots showing the distribution of values for PEER factor 1 ‘extrinsic’ **(A)** and PEER factor 9 ‘intrinsic’ **(B)** The effect of conditions (Fn1, red; Fn5, green; Fn25, blue;) is visible. The genetic concordance between two clones of cells from the same donors is shown for both factors **(C,D)** and the loading of phenotypic features is detailed **(D,E)**.

To evaluate whether any PEER factor(s) captured intrinsic variance, we compared their values in cases where more than one cell line from the same donor was available. PEER factor 9 showed no effect of fibronectin concentration (Figure 3B) but the highest genetic concordance (Figure 2D). This factor (which we named the ‘intrinsic factor’) accounted for 5% of the total variance (Supplementary Figure 3). Phenotypic features describing EdU labelling and other nuclear properties, both in single and clumped cells, loaded onto the intrinsic factor (Figure 3F).

### Identification of genes correlating with extrinsic and intrinsic variation

To identify genes whose expression could contribute to phenotypic variance we performed a correlation analysis between the intrinsic and extrinsic factors and gene expression array data independently generated from cell pellets as part of the HipSci resource^7^. Expression of 4573 genes correlated with either the extrinsic or intrinsic factor or both, in at least one fibronectin concentration. From this list, we filtered out genes that were not associated with any Ensembl identifiers. We also removed genes for which multiple probes showed opposite correlation values. The resulting dataset consisted of 3879 genes (Supplementary Table 2), 1321 correlating with the extrinsic factor, 1977 correlating with Intrinsic Factor and 581 with both (Figure 4A).

**Figure 4.**
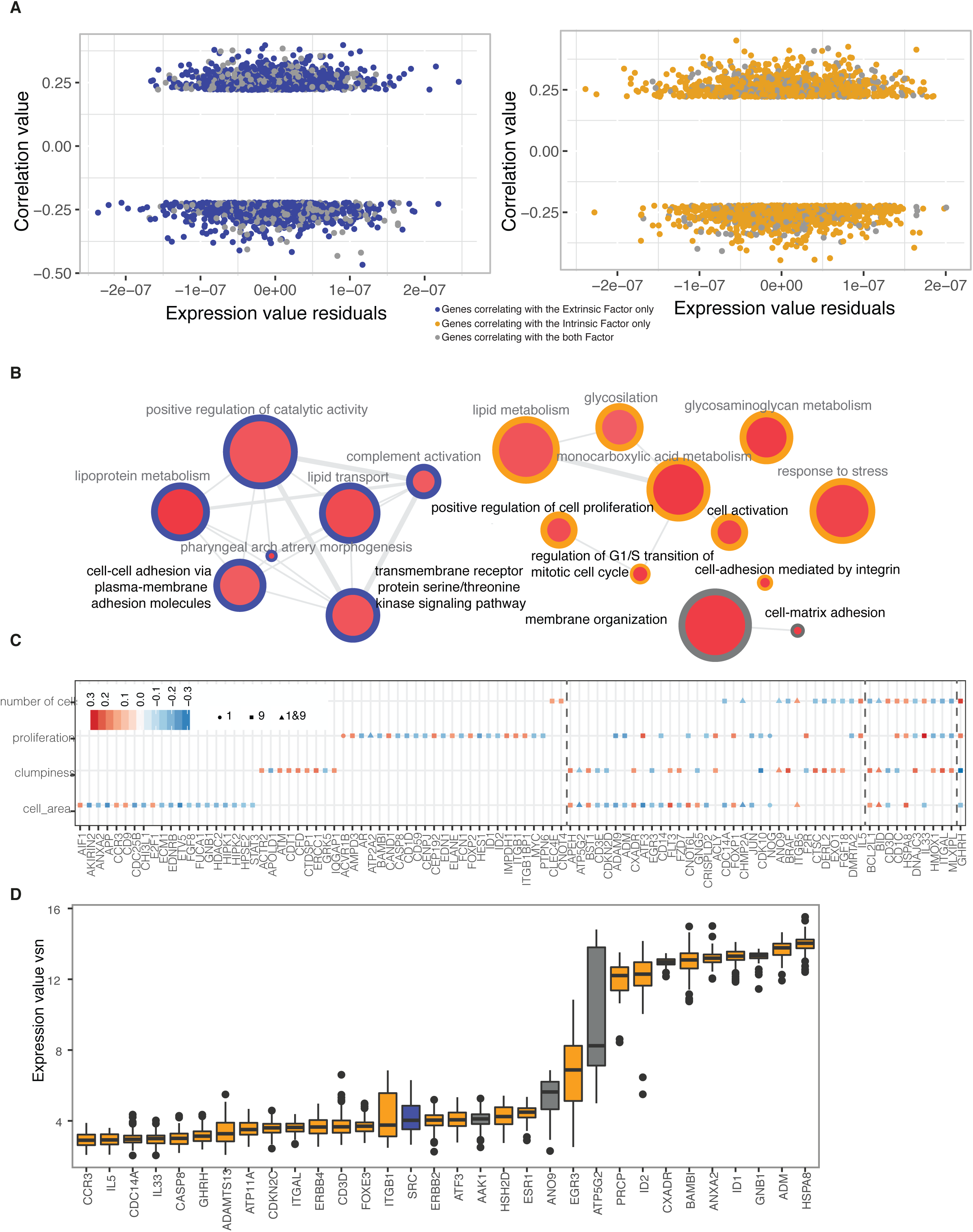
Using the ‘extrinsic’ and ‘intrinsic’ factors to identify genes associated with specific outlier phenotypes. **(A)** A total of 3879 genes correlated with either the ‘extrinsic’ PEER factor (a, blue spots), the ‘intrinsic’ factor (b, yellow spots) or both (a, b, grey spots). (**B)** GO analysis of genes correlating with either or both PEER factors. Circle colours as in (a). Node size (red fill) indicates the p-value associated to each term. Each gene was mapped to the most specific terms applicable in each Ontology. Highly similar GO terms are linked by edges, edge width depicting the degree of similarity. Terms with a black description have been used to select the list of 175 genes. (C) In a total of 98 genes, gene expression correlated significantly with cell area, tendency to form clumps (‘dumpiness’), number of cells and/or proliferation. The colours of the points correspond to the correlation values, while the shapes indicate correlation of a specific gene to the extrinsic and/or intrinsic factors. Grey dotted vertical lines separate genes correlating with one, two or four phenotypes (left to right). **(D)** Boxplots showing the expression values (vsn) of 32 out of the 38 genes with outlier gene expression in one or more outlier cell lines. Color code as in (a).

GO analysis was performed on genes correlating with either or both PEER factors at a threshold value of ± 0.2 of the correlation coefficient (Figure 4B). GO terms associated with the extrinsic factor included cell adhesion and receptor serine/threonine kinase signalling. Terms associated with the intrinsic factor included cell proliferation, response to stress and integrin-mediated cell adhesion. Only two GO terms were associated with both intrinsic and extrinsic factors: membrane organisation and cell-matrix adhesion.

Based on the phenotypes measured in our study, we further filtered the genes according to the functions of their protein products. We selected 175 genes we reasoned were likely to be involved in the distinct cell behaviours observed, belonging to six GO categories describing cell-cell adhesion, regulation of cell cycle and regulation of cell proliferation, cell matrix adhesion, membrane organization and transmembrane receptor signalling pathways. The expression of 98 out of the 175 genes showed a statistically significant correlation with the original phenotypic measurements depicting cell morphology, adhesion and proliferation.

Examples of gene expression variation among cell lines for genes correlating with one, two, three and four phenotypic features are shown in Figure 4C. We noted that most genes showed distinct correlations with the intrinsic and extrinsic PEER factors (Fig. 4C). In addition, opposite correlations were found for a given gene and one or more phenotypes. For example, ITGAL, which mediates intercellular adhesion, was positively correlated with clumping and negatively correlated with proliferation.

38 out of 175 genes showed outlier expression in one or more cell lines (5^th^ and 95^th^ percentiles) (Figure 4D). Almost all of them (32 out of 38) were outliers in outlier cell lines (Figure 4E and Supplementary Table 5). The only outlier gene exclusively associated with the extrinsic PEER factor was SRC, Proto-oncogene tyrosine-protein kinase. In cases in which two cell lines from the same donor were outliers for the same raw phenotypic features (Figure 2), this could not be explained by the overexpression or lack of expression of the same set of genes.

In conclusion, we could identify a set of genes known to regulate cell phenotypic features and observed a correlation between mRNA expression in cell pellets and cell phenotypic features on three concentrations of Fibronectin. Nonetheless, the expression of specific mRNAs was not correlated with outlier cell behaviour.

### Identification of SNVs in cell adhesion genes that correlate with outlier cell phenotypes

To complement the analysis of gene expression variation, we explored whether the presence of Single Nucleotide Variants (SNVs) in gene exons affecting protein function could account for outlier cell behaviour. We observed that donors for which one line was not selected further in the HipSci project for genomic analysis tended to show less genetic concordance (see Figure 2D).

We searched for SNVs in the 3879 genes identified with the extrinsic and intrinsic PEER factors (Supplementary Table 6). Of the 9831 SNVs identified, 4339 were classified as rare, based first on the 1000 Genomes Project ^24^ and ExAC^25^, and second on the frequency in our cell lines (present in fewer than 5 out of 110 lines, Figure 5A). We further filtered the SNVs using the computational model DUET^26^ to predict SNVs that would be deleterious and the computational model Condel^27^ to predict a final list of 95 rare, deleterious and destabilising SNVs that would impair protein structure. The genes that we identified (Supplementary Table 4) encoded proteins that were associated with cell adhesion, including integrins, cytoskeleton and ECM proteins.

**Figure 5.**
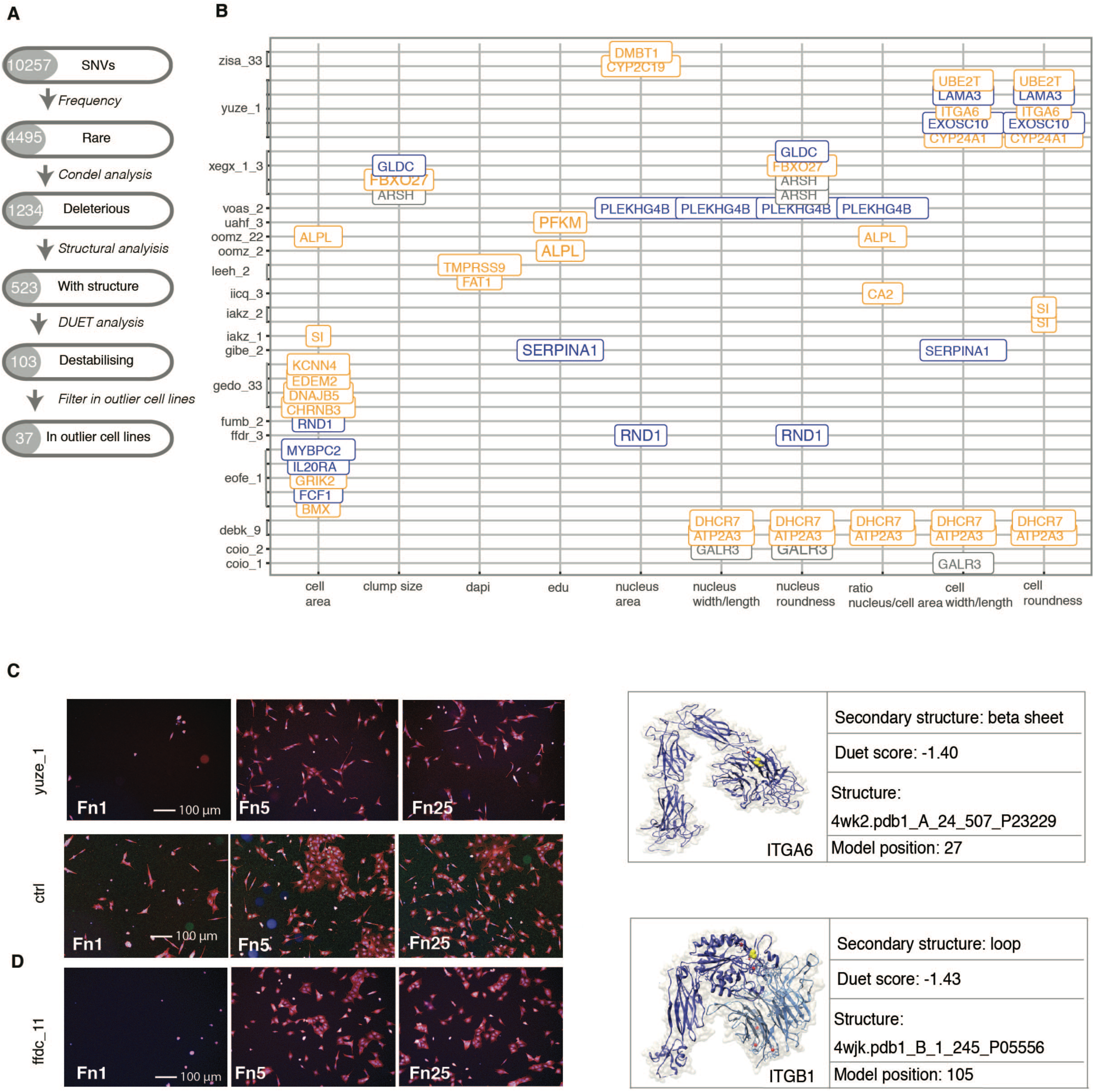
**(A)** Analysis pipeline for selection of genes. (**B)** Genes with at least one rare, deleterious and destabilizing SNV in at least one cell line (y-axis) found to be an outlier for one or more phenotype (x-axis). See Figure 2 for outlier KS analysis. **(C)** Panels show representative images of outlier cell line yuze1, with SNP in ITGA6 control cell line Bob and line ffdc11 harbouring a SNP in ITGB1, right. ITGA6 protein structure depicting position of disruptive SNV in extended beta-strand in yuze_1. Panels show representative images of yuze_1 cells (bottom) and cells from control line BOB (top) plated on different fibronectin concentrations (Fn1, Fn5, Fn25). **(D)** Evaluation of a cell lines not analysed in the original screen (ffdc_11) harbouring a potentially disruptive SNVs in ITGB1 gene. Panels show representative images of cells plated on different fibronectin concentrations (Fn1, Fn5, Fn25).

33 out of 95 rare, deleterious and destabilising SNVs identified occurred in cell lines that were outliers for one or more phenotypes (Figure 5B and Figure supplement 5). An interesting example is the presence of deleterious and destabilising SNVs in the integrin ITGA6 (position 27) in the cell line yuze_1, detected as an outlier for cell and nucleus ratio width/length and for EdU and DAPI intensity. These cells showed reduced spreading (enhanced cell roundness) particularly on the lowest concentration of Fibronectin (Figure 5C and Supplementary Table 4).

Having acquired cell behaviour data for 110 of the HipSci lines, we next explored the genome sequences of the full collection of over 700 lines in the HipSci resource to test whether the presence of rare, deleterious and destabilising SNVs could correctly predict outlier cell behaviour. We identified a SNV in the integrin ITGB1 in the cell line ffdc_11, which like the ITGA6 SNV in yuze_1, maps to the ligand binding domain. When plated on Fibronectin, the ffdc_11 line exhibited reduced attachment and spreading on the lowest Fibronectin concentration (Figure 5C), indicative of the predicted outlier phenotype.

## Discussion

Genetic mapping provides an unbiased approach to discovering genes that influence disease traits and responses to environmental stimuli such as drug exposure^30^. The attractions of developing human in vitro models that reflect in vivo genetics and physiology for mechanistic studies are obvious, and include quantitation via high content image analysis and the replacement of animal experiments. The concept that human disease-causing mutations result in alterations in cell behaviour that can be detected in culture is well established, as in the case of keratin mutations affecting the properties of cultured epidermal cells^31^. In addition, human lymphoblastoid cell lines have long been used to model genotype-phenotype relationships in healthy individuals, although limitations include the confounding effects of biological noise, differentiation state as these cells do not self-renew in vitro and in vitro artefacts such as variation in passage number and growth rate^8,9^.

There has been renewed interest in applying human iPSC for pharmacogenomics, disease modelling and uncovering genetic modifiers of complex disease traits^32,33^. For example, studies with iPS cell derived neurons^34^ support the ‘watershed model’^35^, whereby many different combinations of malfunctioning genes disrupt a few essential pathways to result in the disease. For these reasons we decided to extend the iPSC approach in an attempt to identify genetic modifiers of cell behaviour in healthy individuals. We have recently reported that in an analysis of over 700 well-characterised human iPSC lines there is an 8–23% genetic contribution to variation in cell behaviour^5^. Our ability to detect this contribution depended on the use of simple, short-term, quantitative assays of cell behaviour; the application of multiple environmental stimuli (different concentrations of fibronectin; single cells versus cell clumps); and homogeneous starting cell populations for the assays. The concept that genetic background contributes to variability of human iPSC is supported by a number of earlier studies^13,36,37^.

In order to identify the nature of the genetic contribution to variation in cell behaviour we developed new computational approaches to integrate genomic, gene expression and cell biology datasets. The first was to apply a dimensionality reduction approach, PEER, to capture variance due to extrinsic contributors (different fibronectin concentrations) and genetic concordance. This revealed a robust correlation between RNA expression and the phenotypic features in a large panel of iPSC lines, with specific RNAs associated with intrinsic or extrinsic factors. Carcamo-Orive *et al*. (2017)^3^ also found that human iPSC lines retain a donor-specific gene expression pattern. However, in that study cells were not exposed to different environmental stimuli.

The majority of human iPS cells we screened responded in the same way to all microenvironmental stimuli. This likely reflects canalisation, the process by which normal organs and tissues are produced even on a background of slight genetic abnormalities^38,39^. However, we did identify cell lines that exhibited outlier behaviour that could not be accounted for by variation in gene expression levels, leading us to hypothesise that outlier phenotypes might instead be attributable to genetic variants. We identified rare SNVs that were predicted to be deleterious and for which protein structural information was available. Most of the SNVs identified by this approach occurred in cell lines that were outliers for one or more phenotypes such as cell spreading. The identification of SNVs in integrin genes is of particular interest, because integrins are highly polymorphic and some of the previously reported SNVs alter adhesive function of cancer cells^40,41^. These SNVs not only affected cells in the pluripotent state, but also altered their ability to undergo ectodermal differentiation in vitro, providing proof of principle that our approach can uncover SNVs with lineage-specific effects.

In conclusion, our platform has been successful in associating specific RNAs with intrinsic or extrinsic factors and discovering SNVs that account for outlier cell behaviour. This represents a major advance in attempts to map normal genetic variation to phenotypic variation.

## Materials and Methods

**Cell line derivation and culture** All HipSci samples were collected from skin biopsies of consented research volunteers recruited from the NIHR Cambridge BioResource (http://www.cambridgebioresource.org.uk). Human iPSC were generated from fibroblasts by transduction with Sendai vectors expressing hOCT3/4, hSOX2, hKLF4, and hc-MYC (CytoTune^TM^, Life Technologies, Cat. no. A1377801). Cells were cultured on irradiated or Mitomycin C-treated mouse embryonic fibroblasts (MEF-CF1) in advanced DMEM (Life technologies, UK) supplemented with 10% Knockout Serum Replacement (KOSR, Life technologies, UK), 2 mM L-glutamine (Life technologies, UK) 0.007% 2-mercaptoethanol (Sigma–Aldrich, UK), 4 ng/mL recombinant Fibroblast Growth Factor-2, and 1% Pen**/**Strep (Life technologies, UK). Pluripotency was assessed based on expression profiling^42^, detection of pluripotency markers in culture and response to differentiation inducing conditions^43^. Established iPSC lines were passaged every 3–4 days approximately at a 1:3 split ratio. The ID numbers and details for each cell line are listed in Supplementary Table 1.

**Mycoplasma testing and STR profiling** For mycoplasma testing 1 ml of conditioned medium was heated for 5min at 95°C. A PCR reaction was set up with the following primers: forward (5’GGGAGCAAACAGGATTAGATACCCT3’); reverse (5’TGCACCATCTGTCACTCTGTTAACCTC3'). PCR products were loaded on a 1% w/v agarose gel, run at 110 V for 30 minutes in TAE buffer and observed with Gel Dox XR+ imaging system (Bio-Rad). To confirm cell line identity, DNA extraction was performed using the DNeasy Blood & Tissue Kit (Qiagen). Confluent cells were dissociated from 6-well plates and lysed in protein K solution; 4μl of 100mg/ml RNase solution (Qiagen) was added and DNA was purified through the provided spin-column and eluted in 150μl. DNA quality was confirmed with nanodrop spectrophotometer (Nanodrop 2000, Thermo scientific) and in 1% agarose gel. DNA samples were sequenced using STR profiling at the Wellcome Trust Sanger Institute.

**Fibronectin adhesion assays** 96-well micro-clear-black tissue culture plates (Greiner cat. No. 655090) were coated with three concentrations of human plasma fibronectin (Corning) in alternating columns in a randomised fashion^17^. Cells were incubated for 8 min with Accutase (Biolegend) to create a single cell suspension. As the cells began to separate and round up, pre-warmed medium containing 10 μΜ Rho-associated protein kinase (ROCK) inhibitor (Y-27632; Enzo Life Sciences) was added and cells were removed from culture wells by gentle pipetting to form a single cell suspension. Cells were then collected by centrifugation, aspirated and resuspended in medium containing 10 μM Rock inhibitor. Cells were counted using a Scepter 2.0 automated cell-counting device (Millipore) and seeded onto the fibronectin-coated 96-well plate using Viaflo (INTEGRA Biosciences) electronic pipettes.

Cell line plating was randomised within rows, with three wells per condition for each line to obviate edge and position effects. One control line (A1ATD-iPSC patient 1)^18^, kindly provided by Tamir Rashid and Ludovic Vallier, was run as internal control in the majority of plates. For each well, 3,000 cells were plated for 24 hours prior to fixation. Paraformaldehyde 8% (PFA, Sigma–Aldrich) was added to an equal volume of medium for a final concentration of 4%, and left at room temperature for 15 min. Cells were labelled with EdU (Click-iT, Life Technologies) 30 minutes before fixation. Fixed cells were blocked and permeabilised with 0.1% v/v Triton X-100 (Sigma–Aldrich), 1% w/v bovine serum albumin (BSA, Sigma–Aldrich) and 3%v/v donkey serum (Sigma–Aldrich) for 20 min at room temperature and stained with DAPI (1 microM final concentration, Life Technologies) and cell mask (1:1000, Life Technologies). EdU was detected according to manufacturer’s instructions, except that the concentration of the azide reagent was reduced by 50%.

Images were acquired using an Operetta (Perkin Elmer) high content device. Border wells were avoided to reduce edge effects. The raw Image data are available in the Image Data Resource <placeholder idr0037?> following on from the previous dataset http://dx.doi.org/10.17867/10000107.

Harmony v4.1 software was used to derive measurements for each cell. Measurements included intensity features (DAPI, EdU), morphology features (cell area, cell roundness, cell width to length ratio, nucleus area, nucleus roundness, nucleus width to length ratio) and context features related with cell adhesion properties (number of cells per clump). Processing quantification and normalisation of data was performed as previously described^17^.

### Dimensionality reduction approach

We applied a Bayesian factor analysis model called PEER^16^ to the phenotype data in each cell line. This approach uses an unsupervised linear model to account for global variance components in the data, and yields a number of factor components that can be used as synthetic phenotype in further analysis. We tested a wide range of parameter settings for the model (the *k* number), controlling the amount of variance explained by it. We ran PEER with the full pre-normalized dataset with the following parameters: K = 9; covariates = cell line, fibronectin and batch; maximum iterations = 10,000.

### Gene expression profiling

Gene expression profiles were measured with Illumina HumanHT-12 v4 Expression BeadChips and processed as described in^5^. Probe intensity estimates were normalised separately for the two cell types using the variance-stabilizing transformation implemented in the R/Bioconductor vsn package^44^. After normalisation, the datasets were limited to the final remapped set of probes (nprobes = 25,604). We refer to this version of the gexarray data by vsn log2 (iPS cell/somatic). Open access gene expression array data are available in the ArrayExpress database (https://www.ebi.ac.uk/arrayexpress/) under accession number https://www.ebi.ac.uk/arrayexpress/experiments/E-MTAB-4057/. PEER analysis was performed taking as input the vsn expression values with the following parameters: K = 36; covariates = cell line and batch; maximum iterations = 10,000. The residual gene expression matrix was used to perform a correlation analysis with both intrinsic/extrinsic factor and raw phenotypes using cor() function in R (method Spearman’s). A p-value threshold < 0.05 was applied to select the statistical significant correlations and the cut-off of the correlation values was set to ± 0.2.

### Gene Ontology analysis

Gene Ontology analysis was performed using the Gorilla web-service (http://cbl-gorilla.cs.technion.ac.il/) and the output has been visualised with ReviGO (http://revigo.irb.hr/). Three analyses were performed separately for the genes correlating with the extrinsic factor, the intrinsic factor and both factors.

### Single Nucleotide Variations (SNVs) analysis

All nsSNVs identified from the “INFO_04_filtered” VCF files from the latest release of the exomeseq data, which have been filtered for higher confidence variants using Impute2, were mapped to protein sequences using ANNOVAR^45^. Those nsSNVs which mapped to genes in our set of genes were selected for further analysis.

Rare nsSNVs were defined as those with a minor allele frequency (MAF) < 0.005 in both the 1000 Genomes Project^24^ and ExAC database^25^. Protein domain boundaries were obtained by scanning UNIPROT ^46^ protein sequences against the PFAM^47^ seed libraries using HMMER^48^. UniProt proteins (with mapped nsSNVs) were assigned resolved protein structures/homologs from the PDB biounit database^49^ using BLAST ^50^. Hits were accepted with a sequence identity > 30% and E-value < 0.001. BLAST searches were carried out using both the entire protein sequences and domain sequences.

For each protein with mapped nsSNVs the structural homolog with the highest identity was chosen as a template for homology modelling. In the case of ties the modelling process was performed using each template. The portion of the template and query sequences relating to a BLAST hit were aligned using T-COFFEE^51^. 10 homology models for each query template alignment were created using the MODELLER software^52^. In each case the model with the lowest zDOPE score^53^ was selected for further analysis. Where models were created using several templates the model with the lowest zDOPE out of all created models was selected for further analysis.

The impact of all nsSNVs were assessed using a primarily sequence-based consensus predictor of deleteriousness, Condel^27^. Where structural information was available, the impact of nsSNVs on protein structural stability was also predicted using DUET^26^.

## Acknowledgements

We are grateful to the Wellcome Trust and MRC for funding through the Human Induced Pluripotent Stem Cell Initiative (WT098503). We also gratefully acknowledge funding from the Department of Health via the National Institute for Health Research comprehensive Biomedical Research Centre award to Guy’s & St Thomas’ National Health Service Foundation Trust in partnership with King’s College London and King’s College Hospital NHS Foundation Trust. AV and NML gratefully acknowledge the support of The Francis Crick Institute, which receives its core funding from Cancer Research UK (FC001110), the UK Medical Research Council (FC001110), and the Wellcome Trust (FC001110). AV and NML were also supported by funding from a Wellcome Trust Investigator Award. We thank members of the Watt lab and Jeremy Green for scientific discussions and Mia Gervasio, Sabrina Munir, Ayaulim Nurgozhina, Fatima Chowdhury, Darrick Hansen, Zuming Tang, Christopher Sibley-Allen and Fran Molina for technical support.

