## Supplementary Materials for "Identifying extrinsic versus intrinsic drivers of variation in cell behaviour in human iPS cell lines from healthy donors" (from another version of this preprint: version 1, 2018-03-20)

a

| | | | |
|---|---|---|---|
| Cell area | Clumped cells | mean | 1 |
| | | standard deviation | 2 |
| | Single cells | mean | 3 |
| | | standard deviation | 4 |
| | All cells | mean | 5 |
| | | standard deviation | 6 |
| Cell ratio w/l | Clumped cells | mean | 7 |
| | | standard deviation | 8 |
| | Single cells | mean | 9 |
| | | standard deviation | 10 |
| | All cells | mean | 11 |
| | | standard deviation | 12 |
| Cell roundness | Clumped cells | mean | 13 |
| | | standard deviation | 14 |
| | Single cells | mean | 15 |
| | | standard deviation | 16 |
| | All cells | mean | 17 |
| | | standard deviation | 18 |
| Nucleus area | Clumped cells | mean | 19 |
| | | standard deviation | 20 |
| | Single cells | mean | 21 |
| | | standard deviation | 22 |
| | All cells | mean | 23 |
| | | standard deviation | 24 |
| Nucleus ratio w/l | Clumped cells | mean | 25 |
| | | standard deviation | 26 |
| | Single cells | mean | 27 |
| | | standard deviation | 28 |
| | All cells | mean | 29 |
| | | standard deviation | 30 |
| Nucleus roundness | Clumped cells | mean | 31 |
| | | standard deviation | 32 |
| | Single cells | mean | 33 |
| | | standard deviation | 34 |
| | All cells | mean | 35 |
| | | standard deviation | 36 |
| DAPI | Clumped cells | mean | 37 |
| | | standard deviation | 38 |
| | Single cells | mean | 39 |
| | | standard deviation | 40 |
| | All cells | mean | 41 |
| | | standard deviation | 42 |
| EdU | Clumped cells | mean | 43 |
| | | standard deviation | 44 |
| | Single cells | mean | 45 |
| | | standard deviation | 46 |
| | All cells | mean | 47 |
| | | standard deviation | 48 |
| Number of cells | Clumped cells | | 49 |
| | Single cells | | 50 |
| | All cells | | 51 |
| Clumps | | | 52 |

a, The 52 phenotypic features quantitated for each cell/well

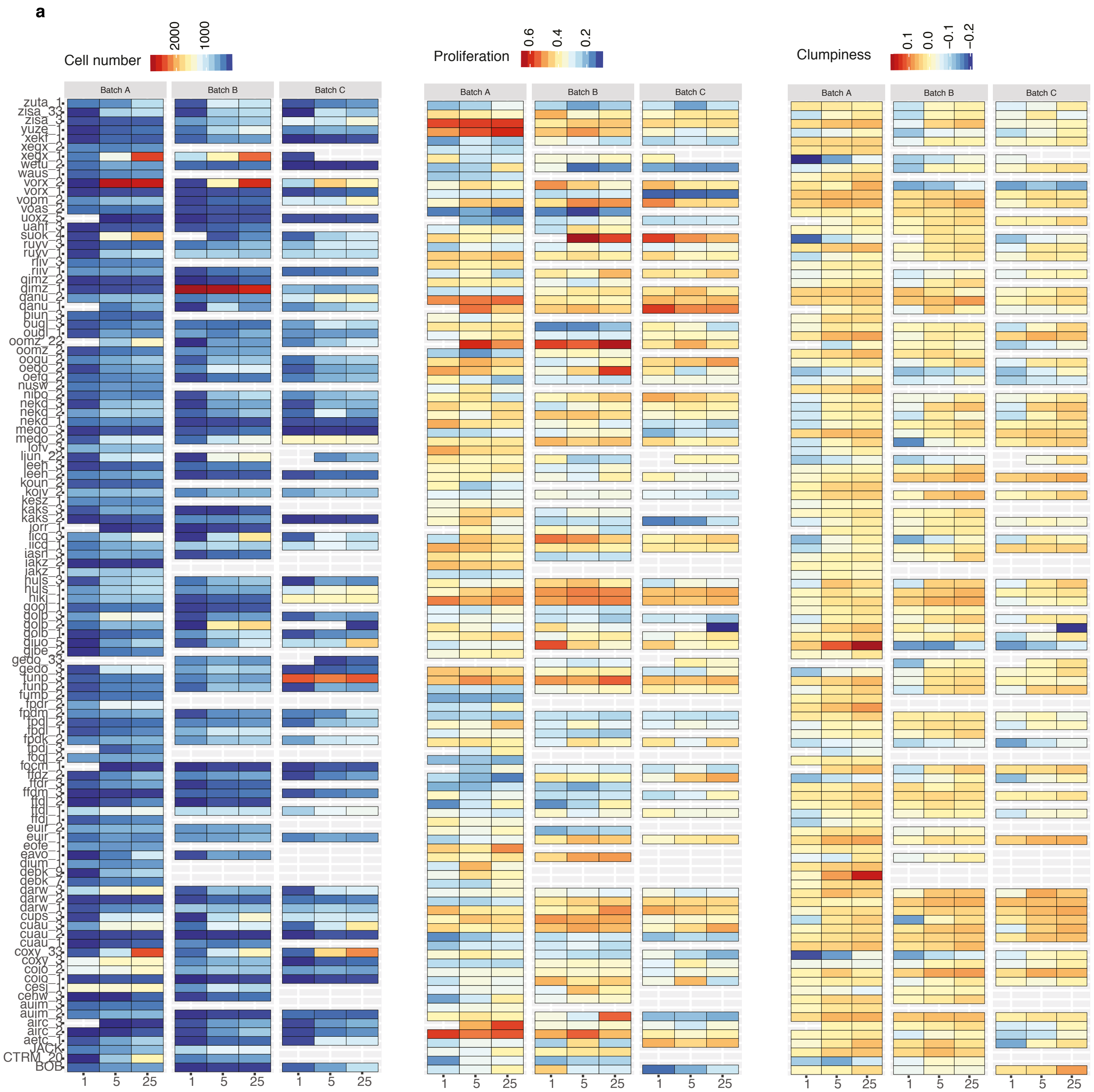


**a,** Heatmap of mean cell number, Proliferation and aggregation measurements for each cell line in three conditions in three biological replicates (batches). Empty boxes correspond to data unavailable or not satisfying the maximum number of cells per well thresholds.

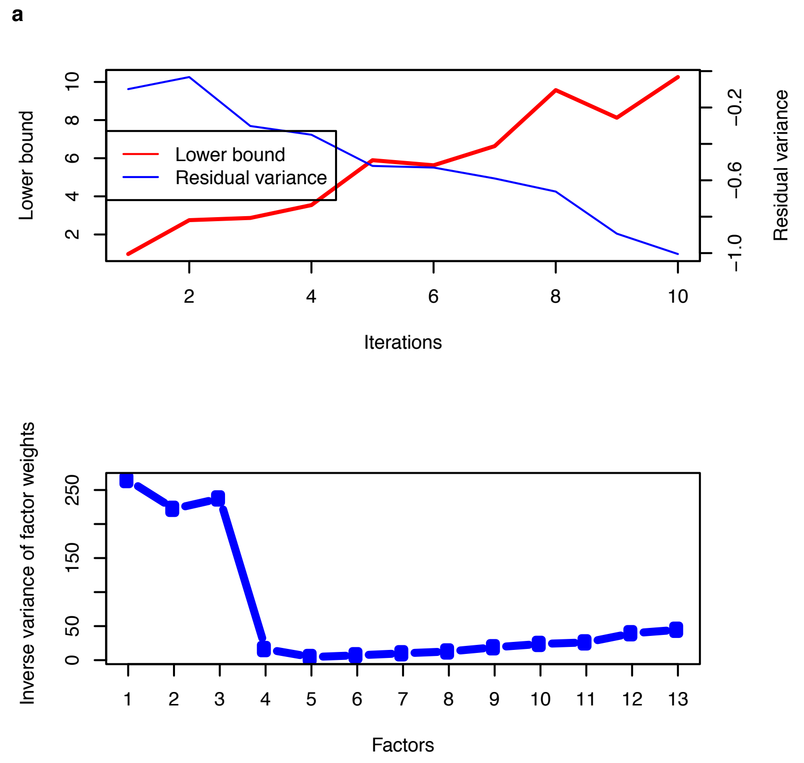


**a,** Automatic relevance detection (ARD) parameters indicates which factors are needed to optimally model the variation in the data.

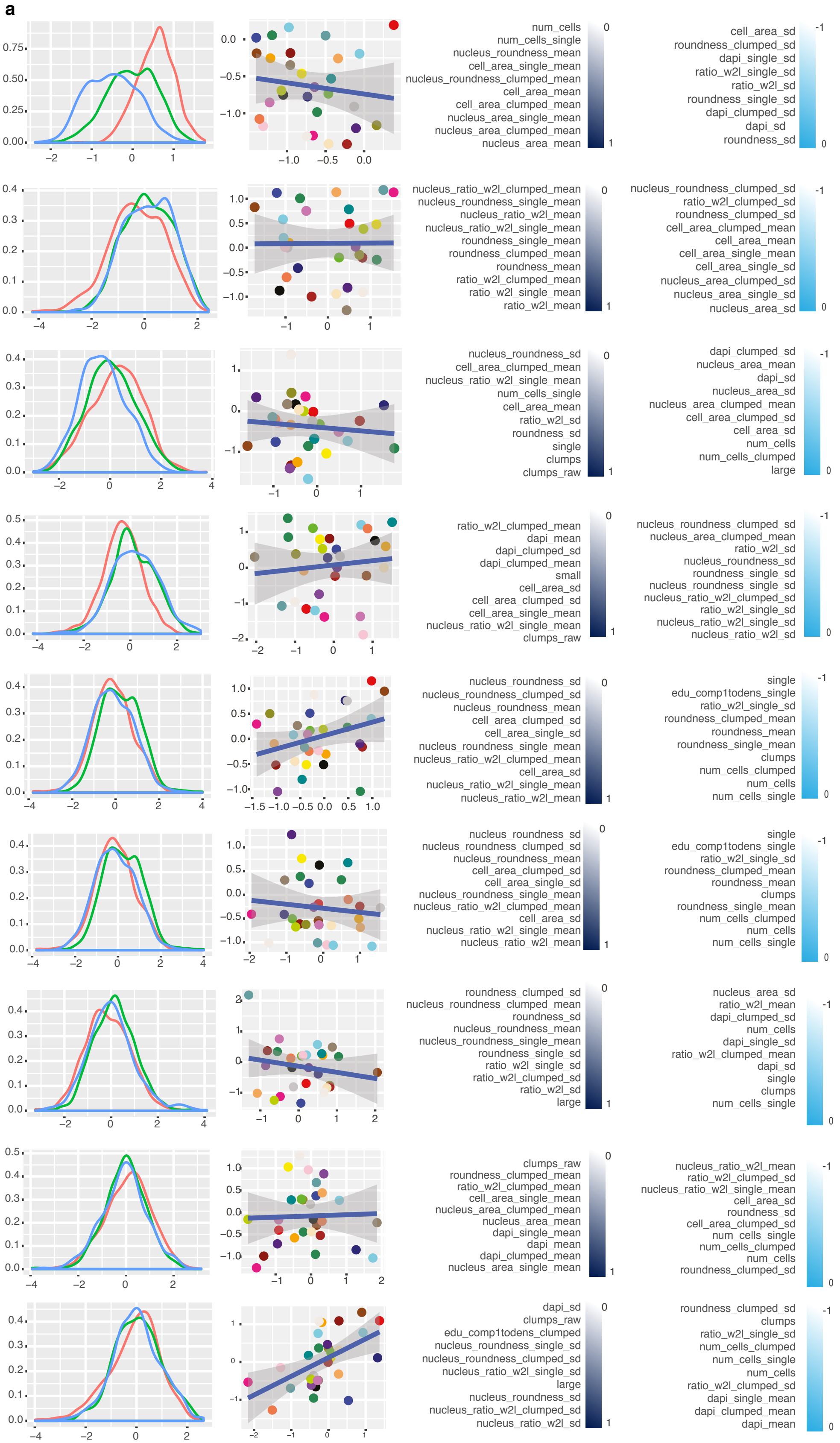


**a,** Nine factors, synthetic phenotypic features obtained with PEER dimensionality reduction. Left, effect of extrinsic conditions (Fn1, red; Fn5, green; Fn25, blue). Middle, genetic concordance between two clones of cells from the same donors. Right, loading of phenotypic raw features into each factor.

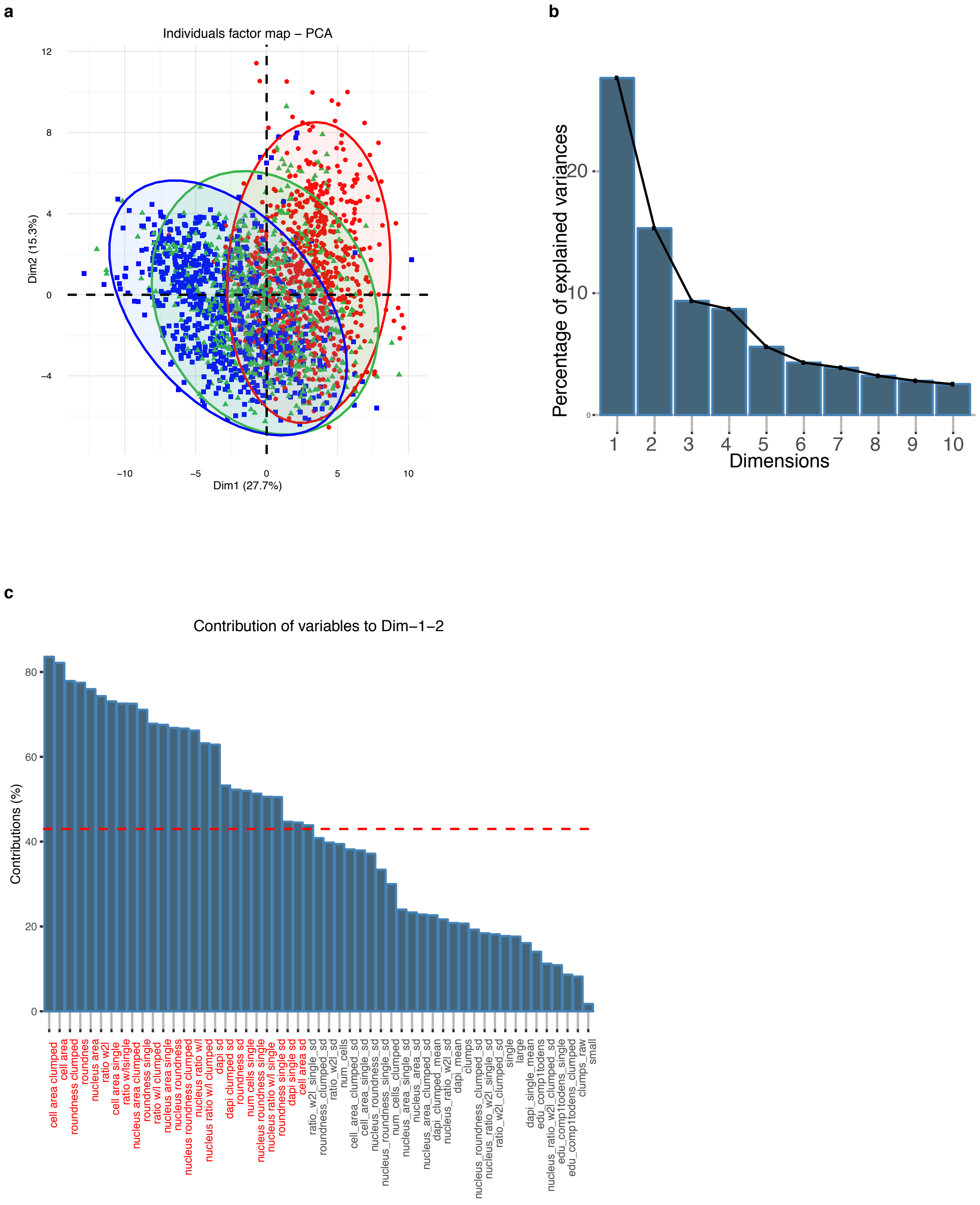


**a,** Principal component analysis is performed on the described phenotypic features. Green, Fn1, red; Fn5, green; Fn25, blue. **b,**The amount of variation retained by each PC.
**c,** The most important variables associated with a given PC 1 and 2. The red dashed line indicates the expected average contribution used here as a cut-off.
Variables with a contribution larger than this cutoff could be considered as important in contributing to the component.

**a**

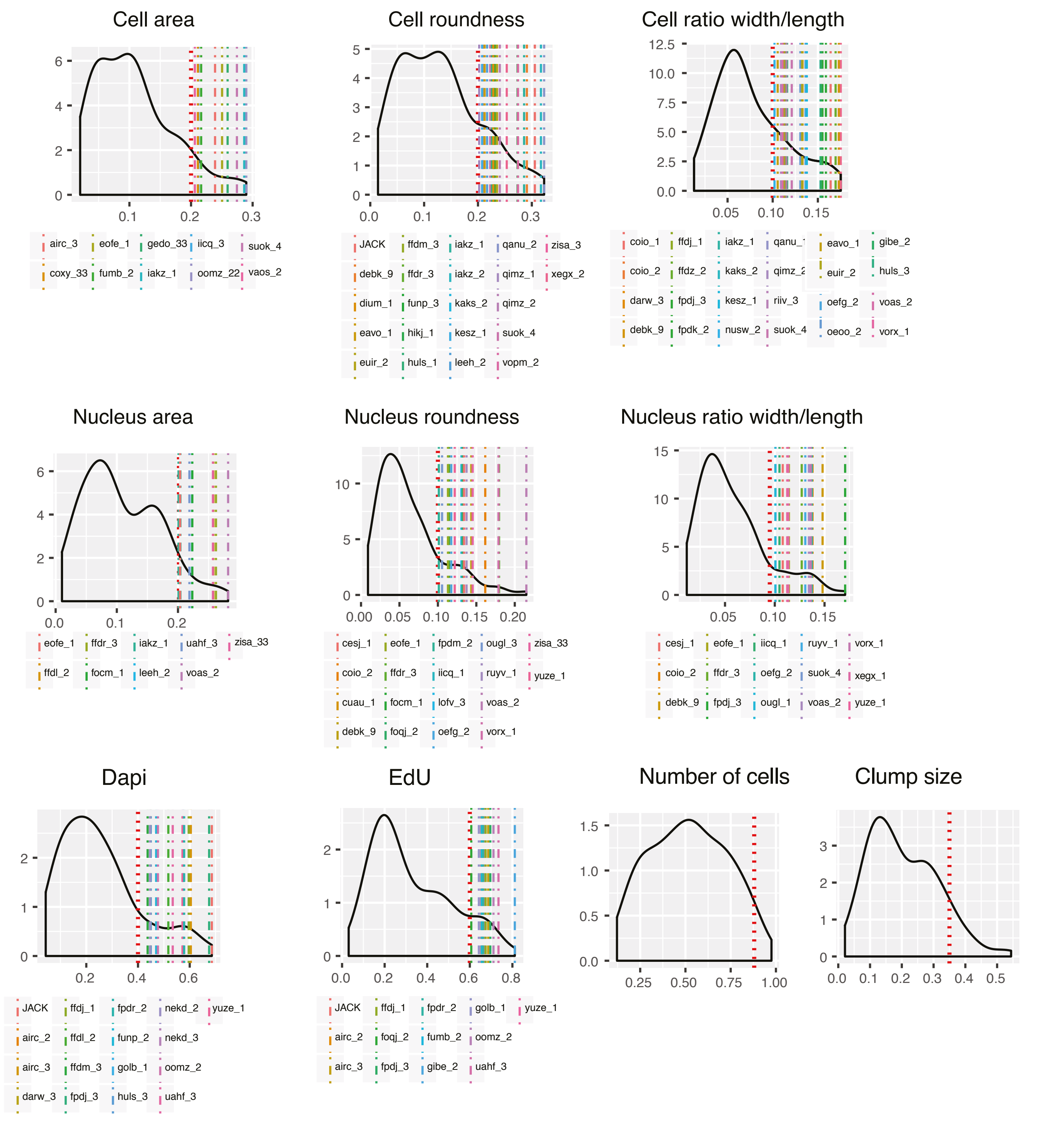


**b**

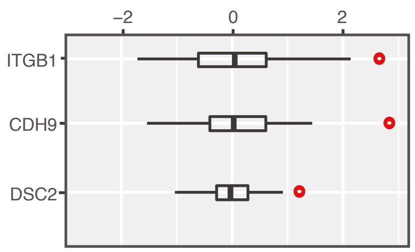


**a,** Distributions of the D scores obtained from ks test for each phenotypic feature. The red dashed line depicts the threshold applyed for each phenotypic feature. The coloured lines represent outlier cell lines. **b,** Variation in gene expression levels.
